# Behavior shapes retinal motion statistics during natural locomotion

**DOI:** 10.1101/2022.09.06.506797

**Authors:** Karl Muller, Jonathan S. Matthis, Kathryn Bonnen, Lawrence K. Cormack, Alexander C. Huk, Mary Hayhoe

## Abstract

Walking through an environment generates retinal motion, which humans rely on to perform a variety of visual tasks. Retinal motion patterns are determined by an interconnected set of factors, including gaze location, gaze stabilization, the structure of the environment, and the walker’s goals. The characteristics of these motion signals have important consequences for neural organization and behavior. However, to date, there are no empirical *in situ* measurements of how combined eye and body movements interact with real 3D environments to shape the statistics of retinal motion signals. Here, we collect measurements of the eyes, the body, and the 3D environment during locomotion. We describe properties of the resulting retinal motion patterns. We explain how these patterns are shaped by gaze location in the world, as well as by behavior, and how they may provide a template for the way motion sensitivity and receptive field properties vary across the visual field.

---

A moving observer travelling through a stationary environment generates a pattern of motion on the retinae that is commonly referred to as optic flow [1], [2]. While optic flow is often thought of as a simple pattern of expansive motion centered on the direction of heading, this is true only in the case of linear motion with gaze centered on heading direction, a condition only rarely met in natural behavior. The actual retinal motion pattern is much more complex, and depends on both the three dimensional structure of the environment and the motion of the eye through space, which in turn depends on the location of the eye in the scene and the gait-induced oscillations of the body. The pervasive presence of self-motion makes it likely that the structure of motion processing systems is shaped by these patterns at both evolutionary and developmental timescales. This makes it important to understand the statistics of the actual motion patterns generated in the context of natural behavior.

Visual motion selectivity is present in the responses of V1 cells, which exhibit both speed and direction selectivity. V1 projects both directly and indirectly to primate area MT [3] which appears to integrate and segment motion signals originating in V1. Additionally, direction encoding in MT is correlated with behavioral variability, suggesting that a motion perception signal upon which decision making is based could originate in MT [4], [5]. Albright [6] observed a centrifugal directionality bias at greater eccentricities in MT, as might be expected from optic flow generated by forward movement, and speculated that this might reflect cortical sensitivity to the statistics of visual motion. However understanding more complex response properties of this and other motion sensitive areas ultimately requires knowledge of the natural statistics that shape neural selectivity. While there has been progress in relating the properties of MT and MST to perception and behavioral goals (see [7] and [8] for reviews), it is difficult to do this effectively without an explicit knowledge of the motion input and how it is shaped by behavior. A similar point was made by Bonnen et al [9] who demonstrated that an understanding of the retinal images resulting from binocular viewing geometry allowed a better understanding of the way that cortical neurons might encode the three-dimensional environment.

Despite many elegant analyses of the way that observer motion generates retinal flow patterns, a detailed understanding has been limited by the difficulties in recording the visual input during locomotion in natural environments. In this paper we examine eye and body movements during locomotion in a variety of natural terrains, and explore how they shape the properties of the retinal input. A number of studies have examined motion patterns generated by cameras moving through natural environments [10], [11], but these data do not accurately reflect the patterns incident on the human retinae. In natural locomotion, walkers gaze at different locations depending on the complexity of the terrain and consequent need to find stable footholds [12]. Thus task goals indirectly affect the motion input. In addition, natural locomotion is not linear. Instead, the head moves through a complex trajectory in space during the gait cycle, while gaze remains stable in the environment, and this imparts a complex pattern of rotation and expansion on the retinal flow [13]. This modulation of the flow pattern with gait raises the question of how the flow patterns are used perceptually, and Matthis et al [13] suggest that walkers learn the time-varying flow patterns that are consistent with postural stability and hence use flow for this purpose. This contrasts with its more familiar role in heading computations. Direct measurement of the motion patterns on the retina during natural locomotion are thus critical for interpreting the perceptual role of motion information, and this requires measurement of eye and body movements during natural locomotion.

In the present work, we simultaneously recorded gaze and image data while subjects walked in a variety of different natural terrains, and reconstructed a 3D representation of the terrain. This links the eye and body movements to the particular terrain and consequently allows calculation of the motion patterns on the retinae. Previous work by Calow and Lappe simulated retinal flow patterns by estimating gaze location and gait oscillations and using a data base of depth images [14, 15]. However, since terrain is a profound influence on gaze deployment, the *in situ* data collection strategy allows a more precise evaluation of the natural statistics than those studies, and we focus on interactions between gaze, body, and the resulting motion patterns in the present work. We find a stereotyped pattern of gaze behavior that emerges due to the constraints of the task, and this pattern of gaze, together with gait-induced head movements, drives much of the variation in the resulting visual motion patterns. Most importantly, walkers stabilize gaze, and the motion statistics result from the motion of the eye in space as it is carried forward by the body while counter-rotating to maintain stability. Because of this stabilization, velocities around the fovea are always low. The periods of stabilization are separated by high velocity saccadic movements to a new gaze point, and the motion generated by these movements must be treated separately because of the mechanisms of saccadic suppression. In all terrains, the ubiquitous presence of the ground plane leads to large speed asymmetries between upper and lower visual fields. As terrain becomes more complex, walkers look closer to the body for foothold information, and this reduces the magnitude of the asymmetry, and also leads to a horizontal band of low velocities centered on the gaze point. Vertical directions predominate in the direction distribution across the visual field, although lateral motion of the body introduces more horizontal components. Thus evaluation of retinal image statistics and their consequences for neural organization requires an understanding of the way the body interacts with the world.

## Results

Eye movements, first person scene video, and body movements were recorded using a Pupil Labs mobile eye tracker and a Motion Shadow full body IMU-based capture system. Eye movements were recorded at 120Hz. The scene camera recorded at 30Hz with 1920×1080 pixel resolution and 100 deg diagonal field of view. The Shadow motion capture system recorded at 100Hz and was used to estimate joint positions and orientations of a full 3D skeleton. Participants walked over a range of terrains 2 times in each direction. Examples of the terrains are shown in Figure 2a. In addition, a representation of the 3D terrain structure was reconstructed from the sequence of video images using photogrammetry, as described below in the section on optic flow estimation. Details of the procedure for calibrating and extracting gaze, body, and terrain data are described in the Methods section, as well as in [12] and [13].

### Oculomotor patterns during locomotion

Figure 1a shows a schematics of the typical eye movement pattern seen in locomotion. When the terrain is complex, subjects mostly direct gaze toward the ground a few steps ahead [12]. This provides visual information to guide upcoming foot placement. As the body moves forward, the subject makes a sequence of saccades to locations further along the direction of travel. Following each saccade, gaze is held approximately stable for periods of 200-300 msec so that visual information about upcoming foothold locations can be acquired while the subject moves forwards during a step. This results in downward slow rotations of the eye to offset the forward motion of the body. Figure 1b shows an excerpt of the vertical component of gaze during this saccade-stabilize gaze pattern. Stabilization is most likely accomplished by the vestibular ocular reflex, although other eye movement systems might also be involved [13]. (See Video demonstrating this behavior [13])

**Figure 1:**
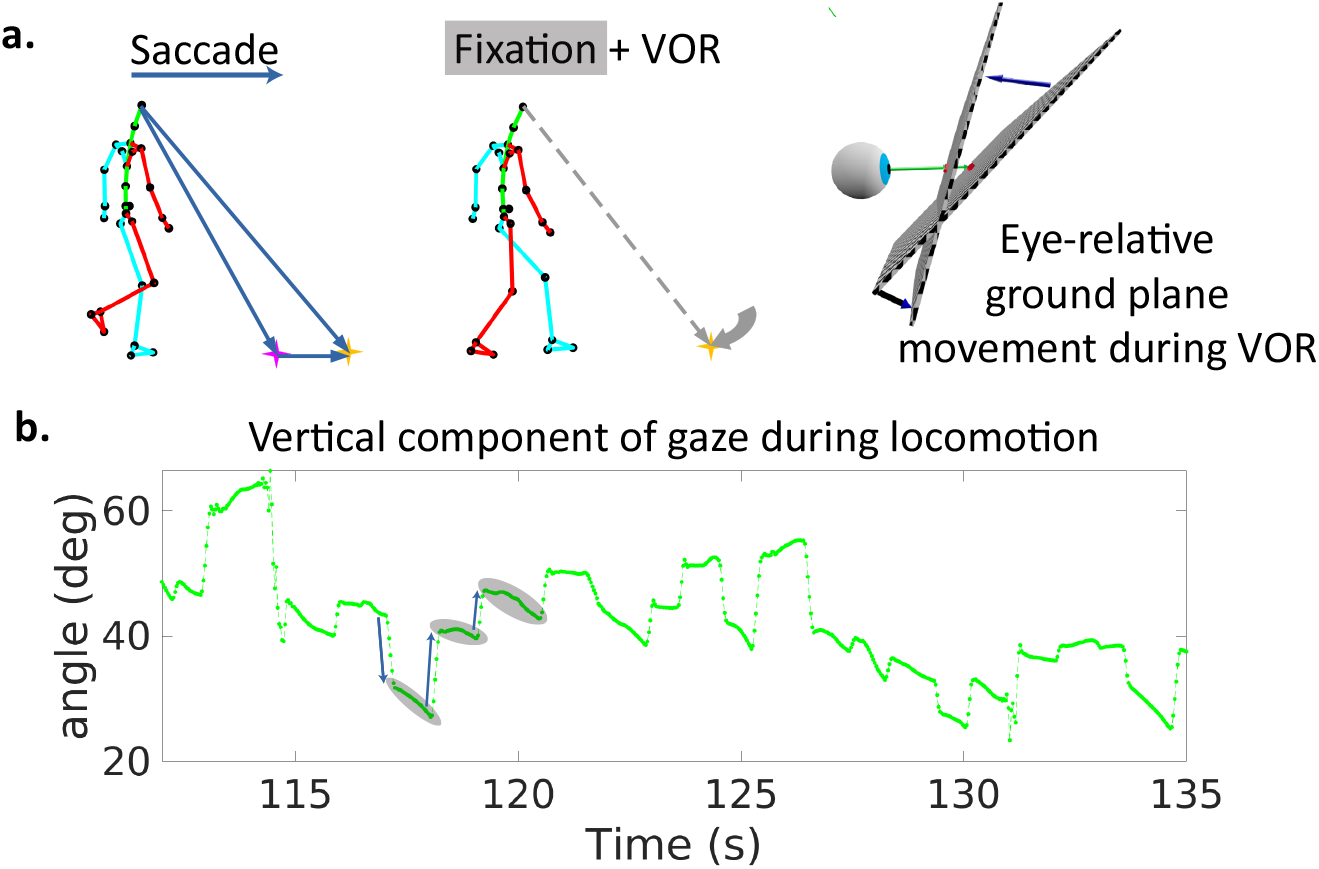
Saccade-stabilization patterns are a common feature of oculomotor behavior during locomotion. **a**. Schematics of saccade and stabilization present in locomotion when looking at the nearby ground. In the top left, the walker makes a saccade to an object further along the path. In the middle panel, the walker fixates this object for a time. The right panel shows the ground becoming more normal to the ground plane during stabilization (e.g., VOR). **b**. Excerpt of vertical gaze angle trace during a period of saccade-stabilize. As participants move forward while looking at the nearby ground, they make sequences of saccades (e.g., dark blue arrows) to new locations, followed by fixations (e.g., gray ellipses) where gaze is held stable at a location in the world while the body moves forward along the direction of travel. The saccades are visible in the trace as high velocity jumps (three examples marked with arrows). These are followed by slower counter-rotations of the eye in the orbit in order to maintain gaze at a fixed location in the scene (three examples marked with gray ovals.)

During the periods of stabilization the retinal image expands and rotates, depending on the direction of the eye in space, carried by the body (See Video from demonstration [13]). It is these motion patterns that we examine here. We segmented the image into saccades and fixations using a velocity threshold of 65*deg/s* and an acceleration threshold of 5*deg/s*^2^. The velocity threshold is quite high in order to accommodate the smooth counter-rotations during stabilization. Saccadic eye movements induce considerably higher velocities, but saccadic suppression and image blur renders this information less useful for locomotor guidance, and the neural mechanisms underlying motion analysis during saccades is not well understood [16]. We consider the retinal motion generated by saccades separately as described in Supplementary Materials.

Imperfections in gaze stabilization during a fixation will add image motion. Analysis of the fixations revealed that stabilization of gaze location in the world was very good (see Figure 8 in the Supplementary Materials). Retinal image slippage during fixations had a mode of 0.26 deg and a median of 0.83 degrees. This image slippage reflects imperfect stabilization and eye tracker noise as well as some small saccades mis-classified as fixations. In order to simplify the analysis, we first ignore image slip during a fixation, and do the analysis as if gaze were fixed at the initial location for the duration of the fixation. In the Supplementary Material we evaluate the impact of this idealization and show that it is modest.

There is variation in how far ahead subjects direct gaze between terrain types, as has been observed previously [12] although the pattern of saccades followed by stabilizing eye movements is conserved. We summarize this behavior by measuring the angle of gaze relative to the vertical, and plot gaze angle distributions for the different terrain types in Figure 2. Consistent with previous observations, gaze is moved closer to the body in the more complex terrains, with the median gaze angle in rocky terrain being approximately 45 deg, about 2-3 steps ahead, and that on pavement being to far distances, a little below the horizontal. Note that the distributions are all quite broad and sensitive to changes in the terrain, such as that between a paved road and a flat dirt path. Subtle changes like this presumably affect variation in the nature of the visual information needed for foot placement. The bimodality of most of the distributions reflects the observation that subjects alternate between near and far viewing, presumably for different purposes (for example, path planning versus foothold finding). These changes in gaze angle, in conjunction with the movements of the head, have an important effect on retinal motion speeds, as will be shown below. Thus motion input indirectly stems from behavioral goals.

**Figure 2:**
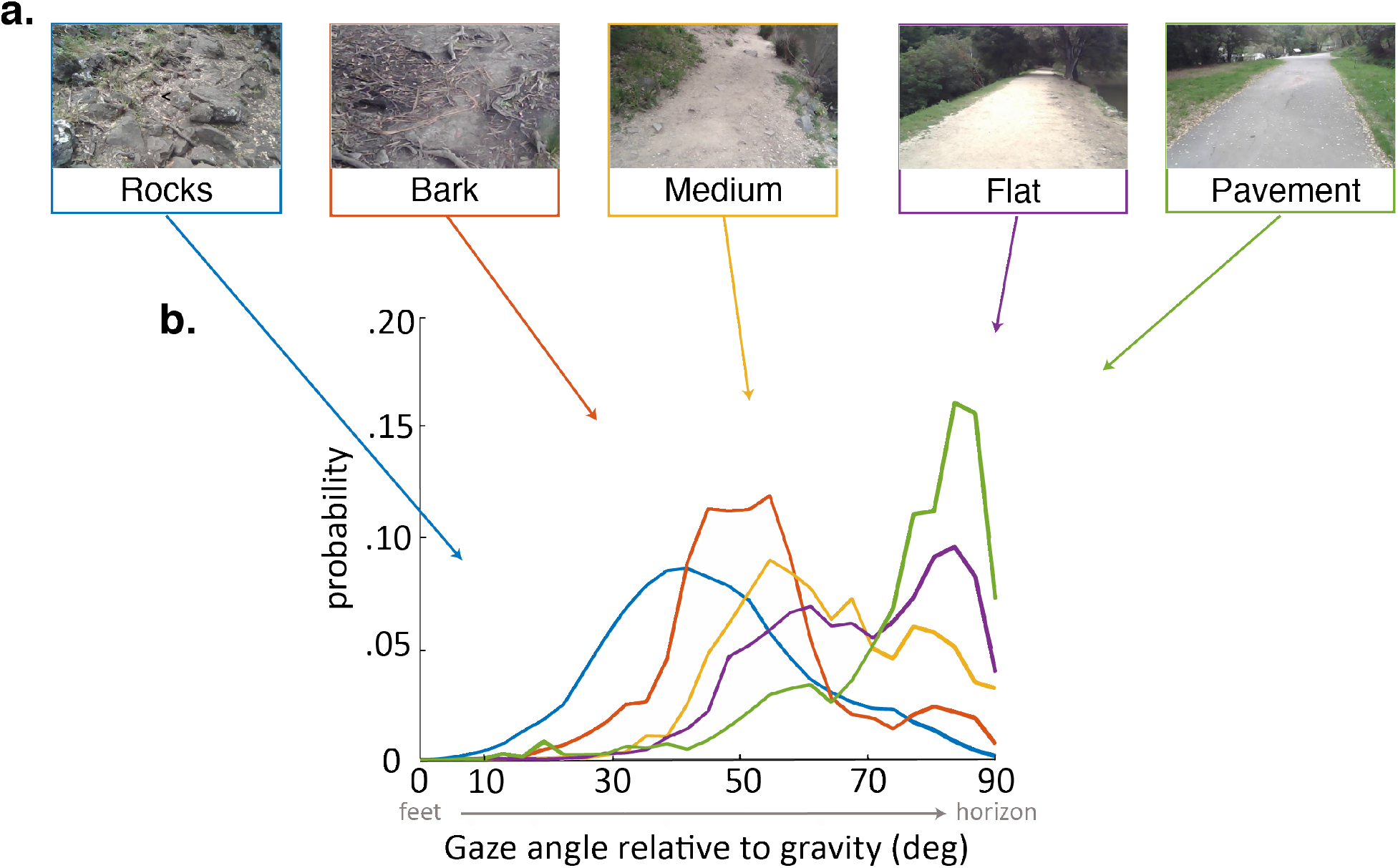
Gaze behavior depends on terrain. **a**: Example images of the five terrain types. Sections of the hiking path were assigned to one of the 5 terrain types. The *Pavement* terrain included the paved parts of the hiking path, while the *Flat* terrain included the parts of the trail which were composed of flat packed earth. The *Medium* terrain had small irregularities in the path as well as loose rocks and pebbles. The *Bark* terrain (though similar to the Medium terrain) was given a separate designation as it was generally flatter than the Medium terrain but large pieces of bark and occasional tree roots were strewn across the path. Finally the *Rocks* terrain had significant path irregularities which required attention to locate stable footholds. **b**: Distributions of vertical gaze angle (angle relative to the direction of gravity) across different terrain types. In very flat, regular terrain (e.g., pavement, flat) participant gaze accumulates at the horizon (90°). With increasing terrain complexity participants shift gaze downwards (30° − 60°).

### Speed and Direction Distributions During Gaze Stabilization

The way the eye moves in space during the fixations, together with gaze location in the scene, jointly determines the retinal motion patterns. Therefore, we summarize the direction and speed of the stabilizing eye movements in Figure 3a-b. Figure 3a show the distribution of movement speeds and Figure 3b shows the distribution of gaze directions (rotations of the eye in the orbit). Rotations are primarily downward as the body moves forward, with rightward and leftward components resulting from both body sway and fixations to the left or right of the future path, occasioned by the need to change direction or navigate around an obstacle. There are a small number of upward eye movements resulting from vertical gait-related motion of the head, and possibly some small saccades that were mis-classified as fixations. These movements, together with head trajectory and the depth structure of the terrain determine the retinal motion.

**Figure 3:**
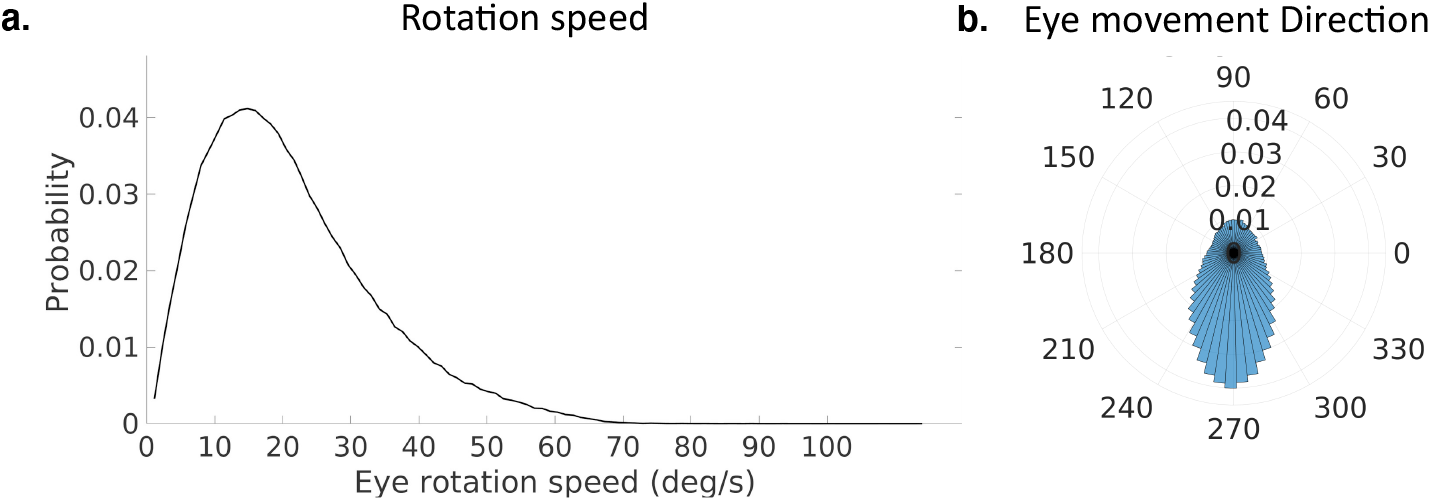
Characteristics of stabilizing eye movements. **a**. The distribution of speeds during periods of stabilization (i.e. eye movements that keep point of gaze approximately stable in the scene). **b**. A polar histogram of eye movement directions during these stabilizing movements. 270 degrees corresponds to straight down in eye centered coordinates, while 90 degrees corresponds to straight up. Stabilizing eye movements are largely in the downward direction, reflecting the forward movement of the body. Some upward eye movements occur and may be due to misclassification of small saccades or variation in head movements relative to the body.

### Optic flow estimation

In order to approximate retinal motion input to the visual system, we first use a photogrammetry package called Meshroom to estimate a 3D triangle mesh representation of the terrain structure, as well as a 3D trajectory through the terrain using the head camera video images. Using Blender [17], the 3D triangle mesh representations of the terrain are combined with the spatially aligned eye position and direction data. A virtual camera is then placed at the eye location and oriented in the same direction as the eye, and a depth image is acquired using Blender’s built in z-buffer method. Thus the depth image input at each frame of the recording is computed. These depth values per location on the virtual imaging surface are mapped to retinal coordinates based on their positions relative to the principal point of the camera. Thus approximate depth at each location in visual space is known. Visual motion in eye coordinates can then be computed by tracking the movement of projections of 3D locations in the environment onto an image plane orthogonal to gaze, resulting from translation and rotation of the eye (see [18] for generalized approach).

The retinal motion signal is represented as a 2D grid where grid points (*x, y*), correspond to polar retinal coordinates (*θ, ϕ*), by the relationship:

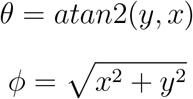

Thus eccentricity in visual angle is mapped linearly to the image plane as a distance from the point of gaze. At each (*x, y*) coordinate there is a corresponding speed in 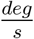 and direction *atan*2(Δ *x*, Δ*y*) of movement.

### Average motion speed and direction statistics

Subjects’ gaze angle modulates the pattern of retinal motion because of the planar structure of the environment [19]. However, we first consider the average motion signal across all the different terrain types and gaze angles. We will then explore the effects of gaze angle and terrain more directly. The mean flow fields for speed and direction, pooled across terrains and subjects are shown in Figure 4. While there will be inevitable differences between subjects caused by the different geometry as a result of different subject heights and idiosyncratic gait patterns, We have chosen to average the data across subjects since the current goal is to describe the general properties of the flow patterns resulting from natural locomotion across a ground plane.

**Figure 4:**
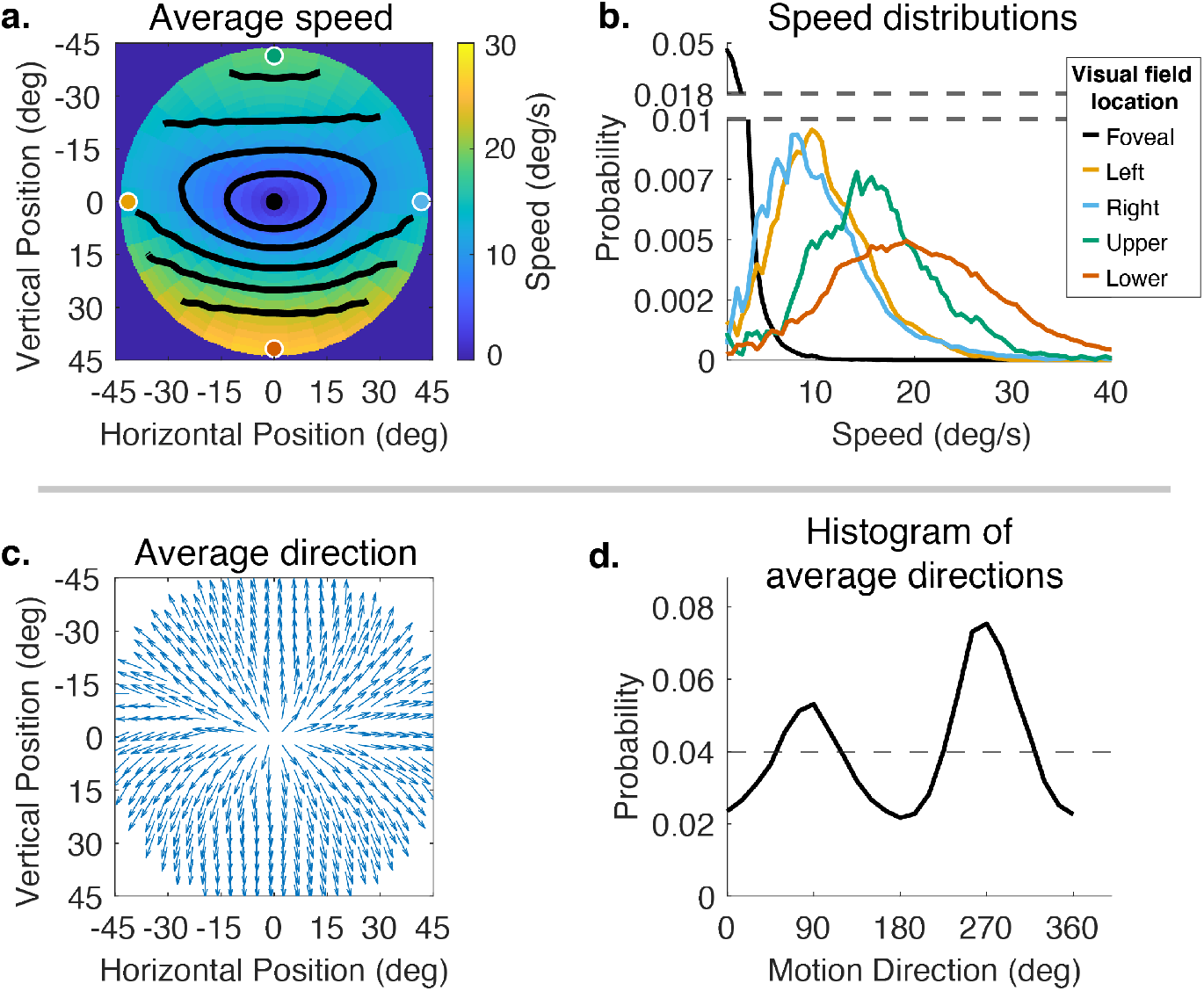
Speed and direction of retinal motion signals as a function of retinal position. **a**. Average speed of retinal motion signals as a function of retinal position. Speed is color-mapped (blue = slow, red = fast). The average is computed across all subjects, and terrain types. Speed is computed in degrees of visual angle per second. **b**. Speed distributions at 5 points in the visual field, at the fovea and four cardinal locations. The modal speed increases in all four cardinal locations, though more prominently in the upper/lower visual fields. Speed variability also increases in the periphery in comparable ways. **c**. Average direction of retinal motion signals as a function of retinal position. Direction is indicated by a unit vector drawn at particular location. Vector direction corresponds to direction in a 2d projection of visual space, where eccentricity from the direction of gaze in degrees is mapped linearly to distance in polar coordinates in the 2d projection plane. **d**. Histogram of the average retinal motion directions (in c.) as a function of polar angle.

Figure 4a shows a map of the average speed at each visual field location (speed is color mapped with blue being the lowest velocity, and yellow being the highest, and the contour lines indicate equal speed). This visualization demonstrates the low speeds near the fovea with increasing speed as a function of eccentricity, a consequence of gaze stabilization. Both the mean and variance of the distributions increase with eccentricity as shown by the speed distributions in Figure 4b. The increase is not radially symmetric. The lower visual field has steeper increase as a function of eccentricity compared to the upper visual field. This is a consequence of the increasing visual angle of the ground plane close to the walker. The left and right visual field speeds are even lower than the upper visual field since the ground plane rotates in depth around a horizontal axis defined by the fixation point (see Figure 2). Average speeds in the lower visual field peak at approximately 20 deg/s (at 45 degrees eccentricity) whereas the upper peaks at 17 deg/s.

Retinal motion directions in Figure 4c are represented by unit vectors. The average directions of flow exhibit a radially expansive pattern as expected from the viewing geometry. However the expansive motion (directly away from center) is not radially symmetric. Directions are biased towards the vertical, with only a narrow band in the left and right visual field exhibiting leftward or rightward motion. This can be seen in the histogram in Figure 4d which peaks at 90 degrees and 270 degrees. Again, this pattern results from a combination of the forward motion, the rotation in depth of the ground plane around the horizontal axis defined by the fixation point, and the increasing visual angle of the ground plane.

### Effects of horizontal and vertical gaze angle on motion patterns

Averaging the data across the different terrains does not accurately reflect the average motion signals a walker might be exposed to in general, as it is weighted by the amount of time the walker spends in different terrains. It also obscures the effect of gaze angle in the different terrains. Similarly, averaging over the gait cycle obscures the effect of the changing angle between the eye and the head direction in space as the body moves laterally during a normal step. We therefore divided the data by gaze angle to reveal the effects of varying horizontal and vertical gaze angle. Vertical gaze angle, the angle of gaze in world coordinates relative to gravity, is driven by different terrain demands that cause the subject to direct gaze closer or further from the body. Vertical gaze angles were binned between 60 and 90deg, and between 17 and 45 deg. This reflects the top and bottom third of the distribution of vertical gaze angles.

The effect of the vertical component of gaze angle can be seen in Figure 5. As gaze is directed more towards the horizontal, the pattern of increasing speed as a function of eccentricity becomes more radially asymmetric, with the peak velocity ranging from less than 5 deg/sec to in the upper visual field to speeds in the range of 20 to 40 deg/sec in the lower visual fields. (Compare top and bottom panels Figure 4a-b.) This pattern may be the most frequent one experienced by walkers to the extent that smooth terrains are most common. As gaze is shifted more towards straight downwards these peaks move closer together and the variance of the distributions increase as the distribution of motion speed becomes more radially symmetric. There is some effect on the spatial pattern of motion direction as well, with the density of downward motion vectors increasing at gaze angles closer to the vertical.

**Figure 5:**
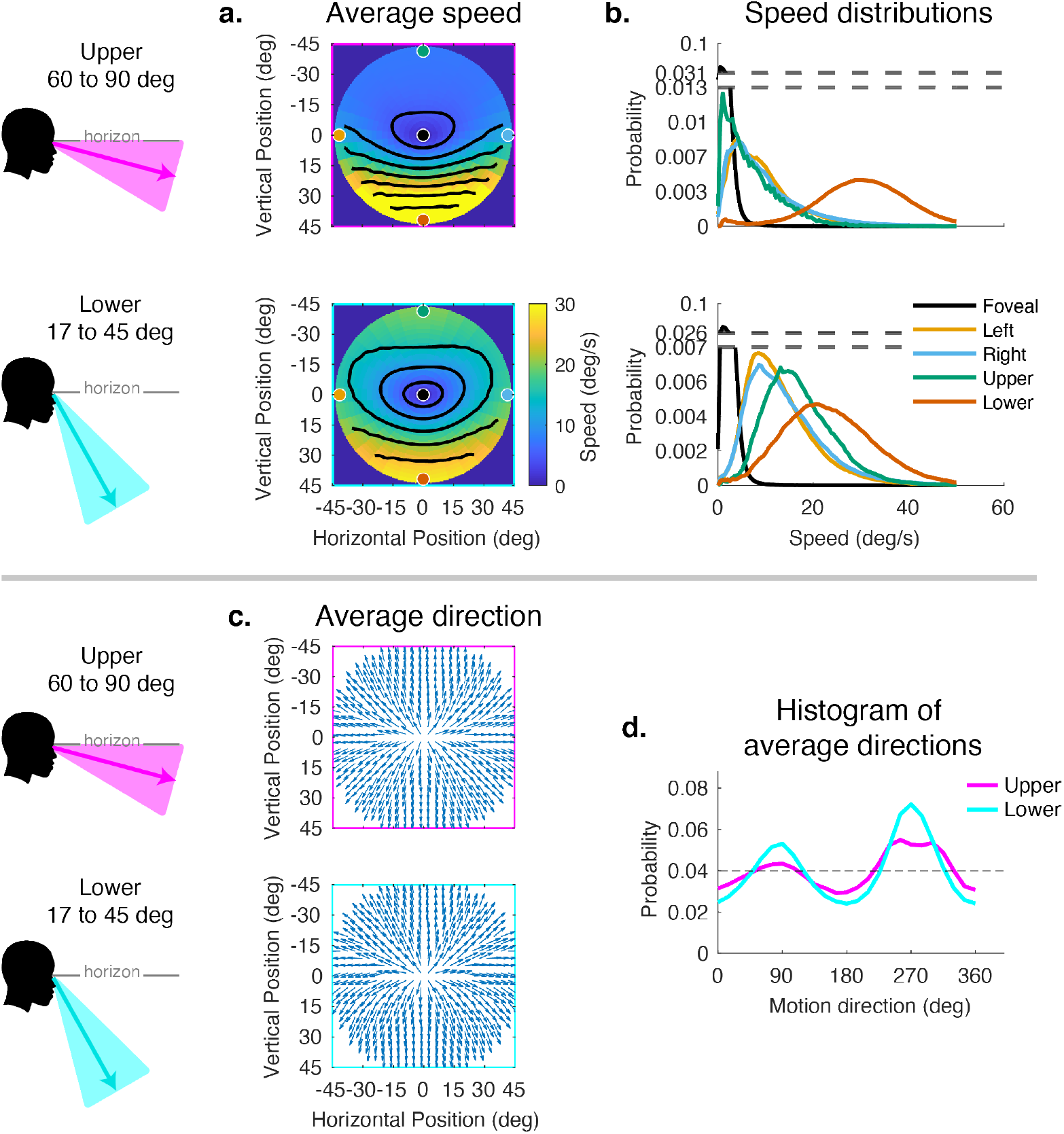
Effect of vertical gaze angle on retinal motion speed and direction. This analysis compares the retinal motion statistics for upper (60°− 90°) vs. lower vertical gaze angles (17°− 45°). The upper vertical gaze angles correspond to far fixations while the lower vertical gaze angles correspond to fixations closer to the body. **a**. Average motion speeds across the visual field. **b**. Five example distributions are shown as in Figure 4. Looking at the ground near the body (i.e. lower vertical gaze angles) reduces the asymmetry between upper and lower visual fields. Peak speeds in the lower visual field are reduced, while speeds are increased in the upper visual field. **c**. Direction maps for upper and lower vertical gaze angles. **d**. Histograms of the average directions plotted in c. While still peaking for vertical directions, the distribution of directions becomes more uniform as walkers look to more distant locations

Horizontal gaze angle is defined relative to the direction of travel. Horizontal gaze angle changes stem both from looks off the path, and from the lateral movement of the body during a step. Body sway accounts for about +/-12 degrees of rotation of the eye in the orbit. Fixations to the right and left of the travel path deviate by about +/-30 degrees of visual angle. Data for all subjects were binned for horizontal gaze angles between -180 to -28 degrees and from +28 to +180 degrees. These bins represent the top and bottom eighths of the distribution of horizontal gaze angles. The effect of these changes can be seen in Figure 6. Changes in speed distributions are shown on the left (Figure 6a-b). The main effect is the tilt of the equal speed contour lines in opposite directions, although speed distributions at the five example locations are not affected very much. Changes in horizontal angle primarily influence the spatial pattern of motion direction. This can be seen in the right side of Figure 6 (in c-d), where rightward or leftward gaze introduces clockwise or counter clockwise rotation in addition to expansion. This makes motion directions more perpendicular to the radial direction of the retinal location of the motion, as gaze becomes more eccentric relative to the translation direction. This corresponds to the curl signal introduced by the lateral sway of the body during locomotion or by fixations off the path [13]. An example of this pattern can be seen in the Matthis et al 2022.

**Figure 6:**
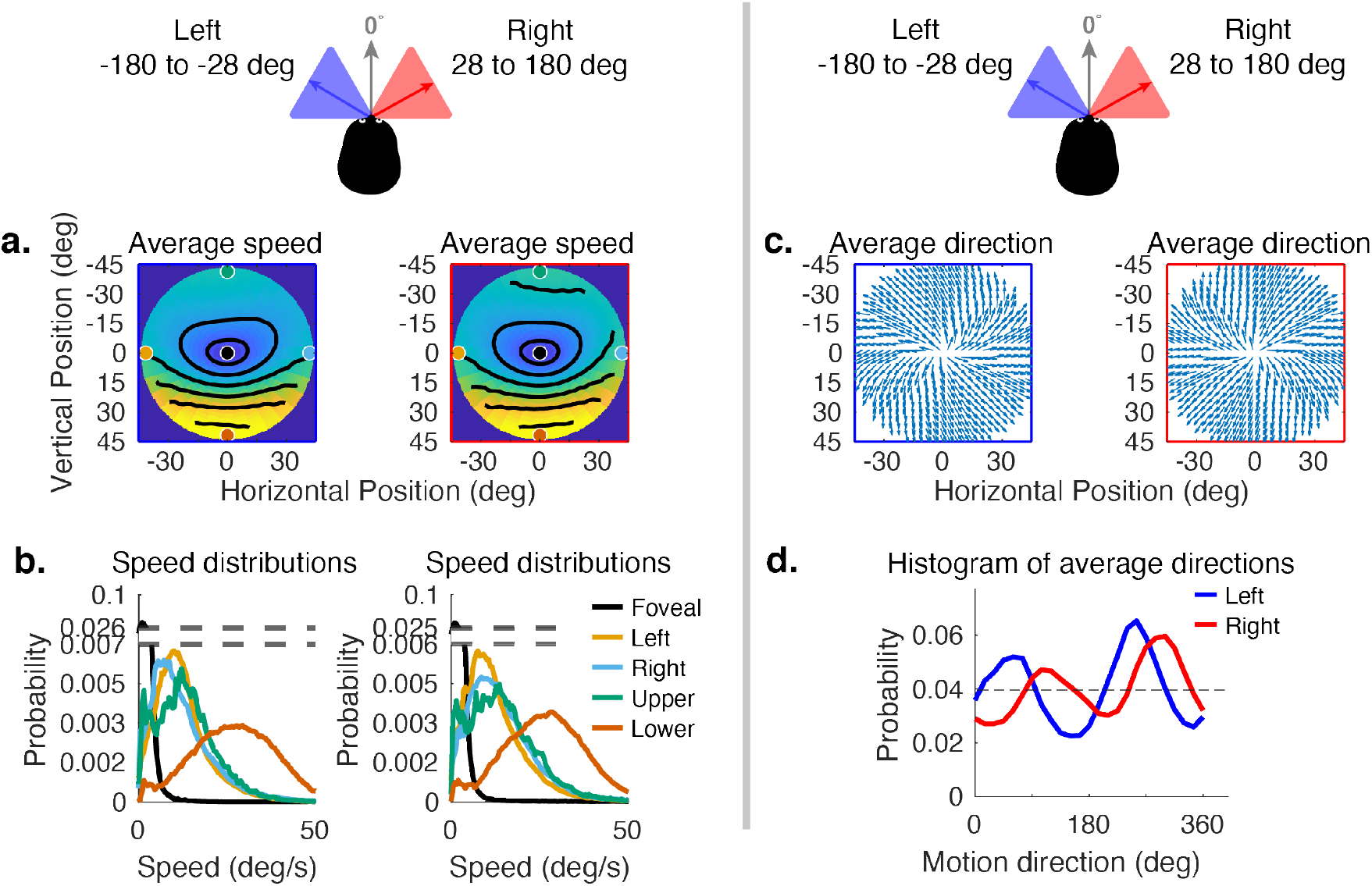
Effect of horizontal gaze angle on motion speed and direction. Horizontal gaze angle is measured relative to the head translation direction. **a**. Average retinal motion speeds across the visual field. **b**. Five distributions sampled at different points in the visual field, as in Figure 4. The effect of horizontal gaze angle on retinal motion directions is unremarkable, except for a slight tilt in to contour lines. **c**. Average direction maps for leftward and rightward gaze angles. **d**. Histograms of the average directions plotted in c. These histograms demonstrate the shift of the rotational component of the flow field.

### Terrain effects on motion

Since subjects generally look close to the body in rough terrain, and to more distant locations in smooth terrain, the data presented thus far confound the effects of 3D terrain structure with the effects of gaze angle. To evaluate the way the terrain itself influenced speed distributions, while controlling for gaze angle, we sampled from the rough and flat terrain data sets so that the distribution of samples across vertical gaze angle were matched. Thus the comparison of flat and rocky terrain reflects only the contribution of the terrain structure to the motion patterns, This is shown in Figure 7. The color maps reveal a somewhat smaller difference between upper and lower visual fields in the rocky terrain than in the flat terrain. This can be seen in the contour plots, and also in the distributions shown on the right. Thus the added motion from the rocky terrain structure attenuated the speed difference between upper and lower visual fields, but otherwise did not affect the distributions very much. There is little effect on the motion direction distributions. This might be expected as the direction and speed of the motion vectors resulting from local depth variations are likely to average out over the course of the path.

**Figure 7:**
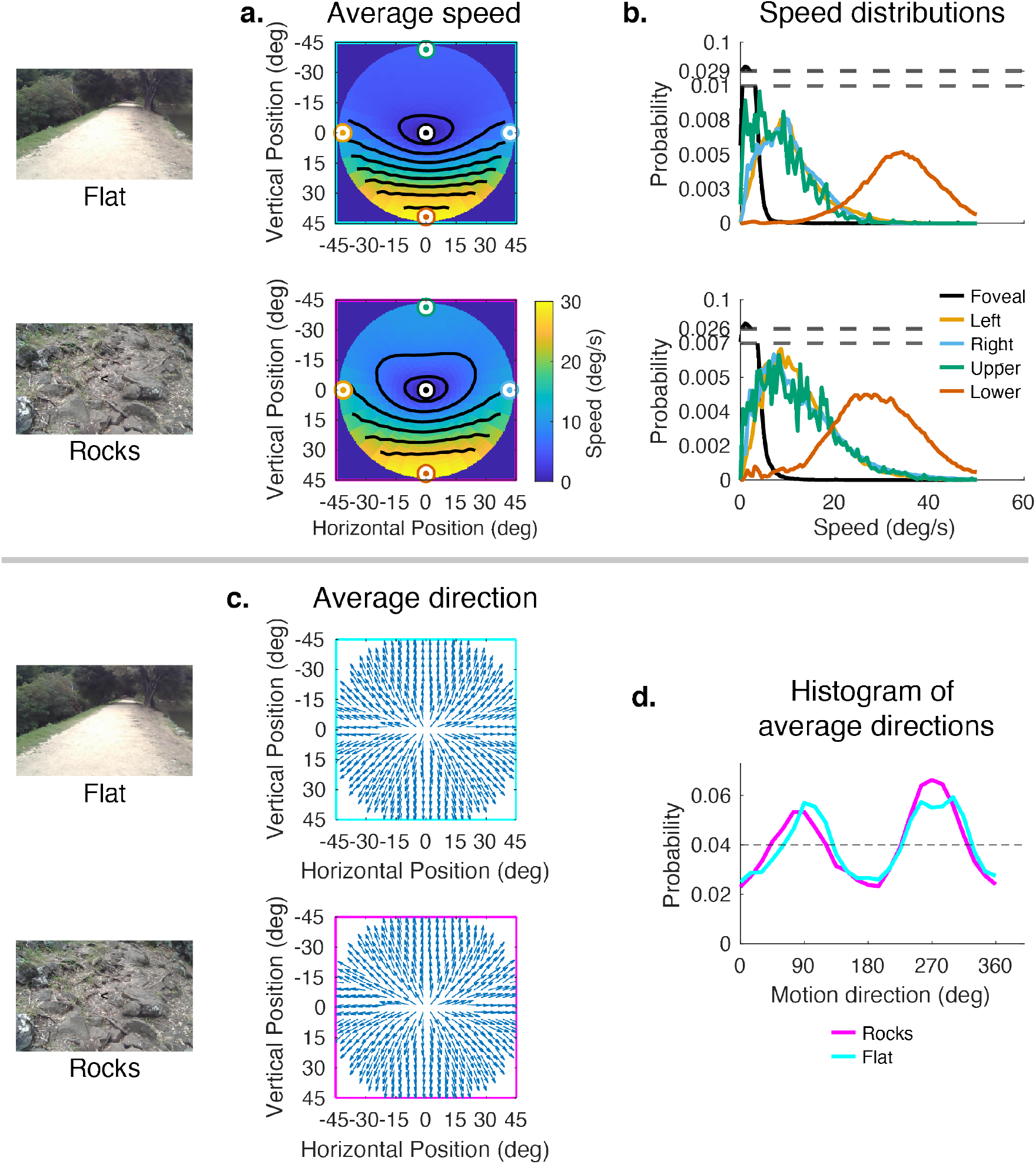
Effect of terrain (independent of vertical gaze) on retinal motion direction and speed. While the vertical field asymmetry is slightly greater for the flat terrain, the effects of terrain on retinal motion direction and speed are modest. **a**. Average retinal motion speeds across the visual field. **b**. Five distributions sampled at different points in the visual field, as in Figure 4. **c**. Average direction maps for leftward and rightward gaze angles. **d**. Histograms of the average directions plotted in c.

## Discussion

We have summarized the retinal motion statistics experienced during locomotion in a variety of natural terrains. This does not sample the broad domain of motion input, most notably self-motion in carpentered environments, where environmental structure may be close to the eye, or environments containing moving objects such as pedestrians and vehicles. However, many of the features of the retinal motion distributions stem from a combination of the ground plane, the saccade and fixate strategy, the gait-induced movements of the head, and the task-linked fixation locations, so one might expect the present observations would cover a substantial domain of experienced motion.

Perhaps the most notable feature of the retinal motion is the ubiquitous pattern of very low velocities at the fovea. This is a consequence the pervasive stabilization of gaze. While the flow field is reminiscent of Gibson’s formulation of optic flow [1], with the focus of expansion centered at the point of gaze, the context is very different from the one Gibson first described, resulting from active stabilization rather than coincidence of gaze with the direction of travel [13]. Calow and Lappe [14, 15] also found generally low velocities at the fovea, but they estimated that the gain of the stabilizing rotations was gaussian with a mean and standard deviation of 0.5. This assumption introduces added motion at the fovea and throughout the visual field. Our measurements indicate that the gain is much closer to 1.0, with only modest retinal slip during fixations (see Supplementary Materials). Given the ubiquitous nature of this kind of stabilizing behavior and the pervasive presence of head movements, the low velocities in the central visual field should be the norm (Land, 2017). Interestingly this result supports the assumption of a motion prior centered on zero, proposed by Wiess et al [20], in order to account for particular motion illusions. Other retinal motion patterns would be experienced during smooth pursuit of a moving target where there would be motion of the background at the fovea. The geometry will also be different for situations that evoke an optokinetic response, or situations where stabilization at the fovea may be less effective, and our results do not pertain to these situations.

While the values of the average statistics computed here reflect the time spent walking in the different terrain types during data collection, rather than the average statistics corresponding to an individual’s experience, the overall pattern reveals features that will generally be true, since they result from walking and fixating at a range of different locations on the ground plane. In particular, the asymmetry between upper and lower visual fields, the low velocities at the fovea, the compression of directions near the horizontal meridian, and the preponderance of vertical directions modulated by lateral gaze positions should all be common features. The asymmetry between upper and lower visual fields and the influence of lateral gaze position was also observed by Calow and Lappe [14, 15]. However, they did not find a relation between retinal speed and direction, whereas we find a band of low speeds in the horizontal direction centered on the fovea. This difference most likely stems from the task-linked fixation locations in the present study. Calow Lappe [14, 15] estimated gaze locations while subjects viewed images of scenes, or walked in different environments. In both cases the fixation distributions were not linked to the depth images in a way that depended on immediate locomotor demands. In particular, the rugged terrains we used demanded fixations lower in the visual field to determine footholds, and this affects the angular rotation of the ground plane in pitch around the axis defined by the fixation position (see Figure 1). The ground fixations also underlie the preponderance of vertical directions across the visual field (see Figure 4). These fixations on the ground close to the body reduce the speed asymmetry between upper and lower visual fields. Added to this, we found that the motion parallax resulting from the rugged terrain structure itself reduces the asymmetry, independently of gaze angle. Thus the behavioral goals and their interaction with body movements and terrain need to be taken into account for a complete description of retinal motion statistics.

The variations of motion speed and direction across the visual field have implications for the interpretation of the behavior of neurons in motion sensitive areas. Callow and Lappe [15] modeled cortical motion processing based on retinal motion signals based on maximizing mutual information the retinal input and neural representations. They found that their model properties were consistent with many of the observed properties of cells in primate motion sensitive areas. Beyeler and colleagues [21] have also argued that selectivity in the motion pathway, in particular, heading direction tuning in MSTd neurons, could simply be an emergent property of efficient coding of the statistics of the input stimulus, without the specific optimization for decoding heading direction. In more recent work, Mineault et al [22] trained an artificial neural network to relate head direction to flow patterns and found units with properties consistent with those of cells in the dorsal visual pathways. Statistical summaries of input signals and the associated behaviors, such as gaze angle, might consequently prove useful in understanding the responses of motion sensitive neurons if response properties arise as a consequence of efficient coding. For example, given the ubiquitous presence of a ground plane, one might expect to find perceptual biases or neural tuning preferences that vary with eccentricity in a manner similar to that in Figure 4. The asymmetry between speed preferences in upper and lower visual fields (as seen in Figure 4) might also be observed in motion sensitive areas as well as other features of the statistics we observe here.

While these theoretical attempts to predict properties of motion sensitive pathways are encouraging, is unclear to what extent known properties of cells in motion sensitive areas are consistent with the observed statistics. Maunsell and van Essen mapped speed preferences of MT neurons as a function of eccentricity out to 24 deg. While there was a weak tendency for preferred speed to increase with eccentricity, the relationship was more variable than expected from the statistics observed here. Comparison of the observed patterns with MST responses is also complex. The role of MSTd in processing optic flow is well studied in both human and monkey cortex [7, 23, 24, 25, 26, 27], and activity in these areas appears to be linked to heading judgments. However, there is no clear consensus on the role of these regions in perception of self-motion. Many of the stimuli used in neurophysiological experiments have been inconsistent with the effects of self-motion, since the region of low or zero velocity was displaced from the fovea. Despite this, neurons respond vigorously to such stimuli. In humans, MST responds to inconsistent motion patterns comparably to consistent patterns [27]. Similarly in monkeys MST neurons respond to both kinds of stimulus patterns [28, 29, 30]. Since a variety of different motion patterns have been used in all these experiments, it may be necessary to simulate the consequences of self-motion with stimuli that are more closely matched to the natural movement patterns of the animal in question, in order to reach more definitive conclusions. In addition, the present work does not consider motion statistics generated by moving objects, during pursuit eye movements or with large nearby objects, and both MT and MST responses may reflect a broader range of stimulus contexts than those considered here.

Analysis of the effect of different horizontal and vertical gaze angles revealed a marked dependence of motion direction on horizontal gaze angle, and of motion speed on vertical gaze angle. Motion directions were biased in the counter clockwise direction when subjects looked to the left of their translation direction, and clockwise when they looked to the right. The distribution of speed became more radially symmetric as subjects looked closer to their feet. These effects have implications for the role of eye position in processing optic flow, given the relationship between eye orientation and flow patterns. This relationship could be learned and exploited by the visual system. Effects of eye position on response properties of visual neurons have been extensively observed in a variety of different regions, including MT and MST [31, 32, 33]. It is possible that such eye direction tuning could be used to compute the expected flow patterns and allow the walker to detect deviations from this pattern. Matthis et al [13] suggested that the variation of the retinal flow patterns through the gait cycle could be learnt and used to monitor whether the variation in body movement was consistent with stable posture. Neurons that compute the consistency between expected and actual retinal motion have been observed in the visual cortex of mice, where firing rate is related to the degree of mismatch between the experimentally controlled and the anticipated motion signal given a particular movement [34]. It is possible that a similar strategy might be employed to detect deviations from expected flow patterns resulting from postural instability.

In summary, we have measured the retinal motion patterns generated when humans walked through a variety of natural terrains. The features of these motion patterns stem from the motion of the body while walkers hold gaze stable at a location in the scene, and the retinal image expands and rotates as the body moves though the gait cycle. The ground plane imparts an increasing speed gradient from upper to lower visual field that is most likely a ubiquitous feature of natural self-generated motion. Gaze location varies with terrain structure as walkers look closer to their bodies to control foot placement in more rugged terrain, and this reduces the asymmetry between upper and lower visual fields, as does the motion resulting from the more complex 3D terrain structure. Lateral rotation of the eye relative to body direction, both during the gait cycle and while searching for suitable paths, changes the distribution of motion directions across the visual field. Thus an understanding of the complex interplay between behavior and environment is necessary for understanding the consequences of self-motion on retinal input.

## Acknowledgments

This work was supported by NIH grants R01 EY05729, T32 LM012414, and U01 NS116377

## Methods

### Experimental Task and Data Acquisition

*Data, as well as analysis and visualization code available upon request*

### Participants

Two datasets were used in this study. Both were collected using the same apparatus, but from two separately conducted studies with similar experimental conditions. One group of participants (n=3, 2 Males, 1 Female) was recruited with informed consent in accordance with the Institutional Review Board at the University of Texas at Austin and collected in an Austin area rocky creek bed. The second participant group (n=8, 4 Males, 4 Females) was recruited with informed consent in accordance with the Institutional Review Board at The University of California Berkeley and collected in a nearby state park.

### Equipment

Infrared eye recordings, first person scene video, and body movements of all participants were recorded using a Pupil Labs mobile eye tracker [35] combined with a Motion Shadow full body IMU based motion capture system (Motion Shadow, Seattle, WA, USA). The eye tracker has two infrared eye cameras, and a single outward facing scene camera. Each eye camera records at 120Hz at 640×480 resolution, while the outward facing scene camera records at 30Hz with 1920×1080 pixel resolution, with a 100 degree diagonal field of view. The scene camera is situated approximately 3cm above the right eye. The Shadow motion capture system is comprised of 17 3-axis accelerometer, gyroscope, and magnetometer sensors. The readings from the suit are processed by software to estimate joint positions and orientations of a full 3D skeleton. The sensors record at 100Hz, and the data is later processed using custom Matlab code (Mathworks, Natick, MA, USA). See [12] and [13] for more details.

### Experimental Task

For both groups of participants (Austin dataset and Berkeley dataset) the experimental task was similar, with variation in the terrain types between the two locations. For the Berkeley participants, the task involved walking back and forth along a loosely-defined hiking trail that varied in terrain features and difficulty. This trail was traversed two times in each direction by each participant. Different portions of the trail were pre-examined and designated as distinct terrain types, being labeled as one of pavement, flat (packed earth), medium (some irregularities such as rocks and roots), and bark (large pieces of bark and roots but otherwise not rocky), and rocks (large rocks that constrained the possible step locations). Examples of the terrain are shown in Figure **??**. For the Austin participants, the task involved walking back and forth along a rocky dried out creek bed. Participants walked 3 times in each direction. This is the same terrain used in [12]. This terrain difficulty is most comparable to the rocks condition in the Berkeley dataset. For each of the rocky terrains, the ground was sufficiently complex that subjects needed to use visual information in order to guide foot placement (see [12] for more details).

### Calibration and post-processing

At the start of each recording session, subjects were instructed to stand on a calibration mat that was used for all subjects. The calibration mat had marked foot locations that were at a fixed distance from a calibration point 1.5 meters away from the foot locations. Subjects were then instructed to maintain fixation on this calibration point throughout a calibration process. The calibration process involved rotating the head while maintaining fixation, to 8 different pre-determined head orientations (the cardinal and oblique directions). This segment of each subjects recording was later used to align and calibrate the data. This was done by finding an optimal rotation that aligned the mobile eye tracker’s coordinate system to that of the motion capture system, such that the distance between the projected gaze vector and the calibration point on the mat was minimized. This rotation was then applied to the position data in the recording, aligning the rest of the recording in space. Each system’s timestamps were then used to align the recording temporally, as timestamps were recorded to a single laptop computer on a backpack worn by subjects throughout the recording. (See [12] for more details). The 100Hz motion capture data was upsampled using linear interpolation to match the 120Hz eye tracker data.

### Fixation detection

During locomotion, walkers typically maintain gaze on a stable location in the world as the body moves forward [12]. During these periods, the eye counter-rotates in the orbit, driven largely by the vestibulo-ocular reflex, although other eye movement systems might also be involved [13]. We refer to these periods of stable gaze in the world in the presence of a smooth compensatory eye rotation as “fixations”, although in more common usage of the term, the eye is stable in the orbit and the head is fixed. In order to analyze the retinal motion input, we needed to differentiate between saccades and periods of stable gaze, since vision is suppressed during saccades, and the visual information used for guidance of locomotion is acquired during the fixation periods. We therefore needed to segment the eye position data into fixations and saccades. The presence of smooth eye movements during stable gaze makes detection of fixations more difficult. We used a velocity and acceleration threshold method with thresholds set such that detected fixations best match hand coded fixations. The velocity threshold was 65*deg/s* velocity and the acceleration threshold was 5*deg/s*^2^. Frames of the recording that fall below these thresholds are labeled as fixation frames, and sequential fixation frames are grouped accordingly into fixation instances. The velocity threshold is quite high in order to accommodate the smooth counter-rotations of the eye in the orbit during stabilization.

### Fixation idealization via initial location pinning

Imperfections in gaze stabilization during a fixation will add image motion. Analysis of fixational eye movements revealed that stabilization of gaze location in the world was very effective. The effectiveness of stabilization was determined by measuring the deviation of gaze from the initially fixated location over the duration of the fixation. This slippage had a median deviation of 0.83 degrees, and a mode of 0.26 degrees (see below for more details). This image slippage reflects imperfect stabilization, small saccades mis-classified as fixations, and eye tracker noise. In order to simplify the analysis, we first ignore image slip during a fixation, and do the analysis as if gaze were fixed at the initial location for the duration of the fixation. We evaluate the impact of this idealization below. Note that in Calow and Lappe’s previous studies they assumed that the stabilization gain varied randomly between zero and 0.5, which would correspond to much less effective stabilization. This estimate was based on experiments with simulated motion (Lappe, 1998), and consequently may reflect different underlying oculomotor mechanisms, given the absence of vestibular input.

After fixations are detected, eye movement directions are computed. This is done by considering sequential frame pairs within a fixation or a saccade. Then eye coordinate basis vectors are calculated for the first frame in a pair. Gaze direction is the third dimension (orthogonal to the plane within which the eye movement direction will be calculated), the first dimension is the normalized cross product between the eye direction and the negative gravity in world coordinates. The second dimension is the cross product between the first and third dimensions. This convention assumes that there is no torsion about the viewing axis of the eye, and so one dimension (the first) stays within a plane perpendicular to gravity. Using these coordinates, the direction of the next eye direction (corresponding to the eye movement occurring over frame pairs) is computed in the reference frame of the first eye direction’s coordinates (*x, y, z*).

### 3D terrain structure measurement using photogrammetry

Optic flow estimation relied on 3D terrain reconstruction provided by Meshroom [36]. Default parameters were used in Meshroom, with the exception of specifying camera intrinsics (focal length in pixels, and viewing angle in degrees). RGB image based scene reconstruction is subject to noise which will be influenced by the movement of the camera and its distance from the terrain. In order to evaluate the accuracy of the 3D reconstruction we took advantage of the terrain meshes calculated from different traversals of the same terrain by an individual subject, and also by the different subjects. Thus for the Austin data set we had 12 traversals (out and back 3 times by 2 subjects.) Easily identifiable features in the environment (e.g. permanent marks on rocks) were used in order to align coordinate systems from each traversal. A set of corresponding points can be used in order to compute a similarity transform between points. Then the iterative closest point (ICP) method is used to align the corresponding point clouds at a finer scale by iteratively rotating and translating the point cloud such that each point moves closer to its nearest neighbor in the target point cloud. The resulting coordinate transformation is then applied to all recordings such that they are all in the same coordinate frame. Foothold locations were estimated using the different aligned meshes, using a method which found the closest location on a mesh to the motion capture system’s foothold location measurement. The distances between corresponding foothold locations between meshes were then measured. There was high agreement between terrain reconstructions, with a median distance of approx 3cm, corresponding to 0.5 degrees of visual angle. Thus the 3D structure obtained is quite consistent between different reconstructions of the same terrain.

## Supplementary Materials

### Retinal Slip During Fixations

To measure the extent of the retinal slip during a fixation, we took the gaze location in the camera image at the first fixation frame, and then used optic flow vectors computed by Deepflow ([37]) to track this initial fixation location for the duration of the fixation. This is done by indexing the optic flow vector field at the initial fixation location, measuring its displacement across the first frame pair, computing its new location in the next frame, and then measuring the flow vector at this new location. This is repeated for each frame in the fixation. The resulting trajectory is that of the initial fixation location in camera image space over the duration of the fixation. For each frame of the fixation, the actual gaze location is then compared to the current location of the initial fixation location (with the first frame excluded since this is how the initial fixation is defined). The locations are converted into 3D vectors as described in, and the median angular distance between gaze location and initial fixation location is computed for each fixation.

Data from this analysis can be seen in Figure 8 which plots the distribution of the retinal slip during fixations. The mode of the distribution is 0.26 deg and median was 0.83 degrees. This is quite good, especially given that the width of a foothold a few steps ahead will subtend about 2 deg and it is unlikely that foot placement requires more precise specification. It is likely that the long tail of the distribution results from errors in specifying the fixations rather than failure of stabilization. In particular, some small saccades are most likely included in the fixations. Other sources of error come from the eye tracker itself. Another source of noise comes from the fact that the image based slip calculations were done at 30Hz. Treadmill studies have found long periods (about 1.5s) of image stability (defined as less than 4 degrees per second of slip) during slow walking, with this being decreased to 213ms for faster walking [38].

**Figure 8:**
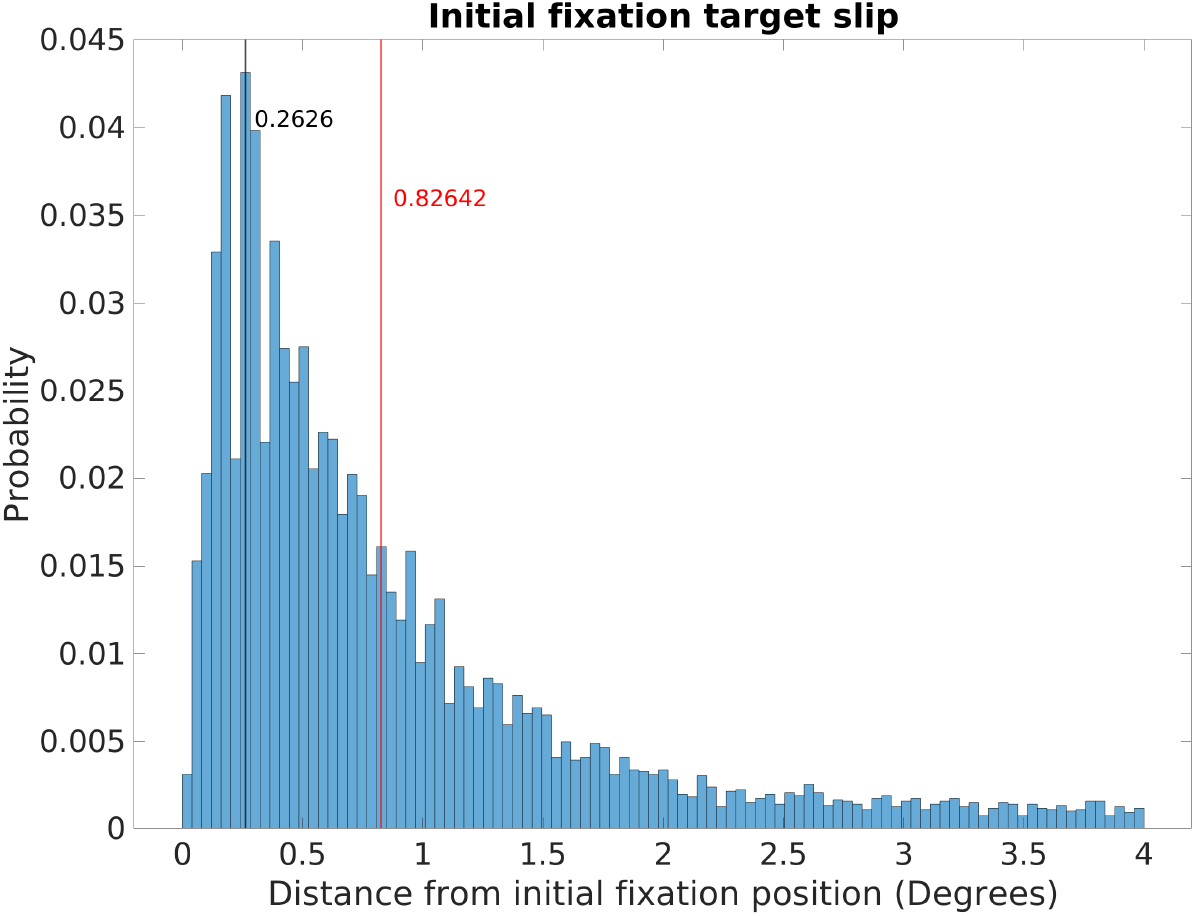
Retinal Image Slippage. Histogram of the deviation from the location of the initial fixation over the course of a fixation. Initial fixation location is computed and tracked over the course of each fixation, and compared to current fixation for duration of each fixation. Median deviation value is calculated for each fixation. The histogram captures the extent of variability of initially fixated locations relative to the measured location of the fovea over the course of fixations, with most initially fixated locations never deviating more than 2 degrees of visual angle during a fixation.

Manual reintroduction of retinal slip (which our measurements suggest arise from a gain in the VOR of <1) simply results in added downward motion to the entire visual field, whose magnitude is equivalent to the slip. This can be seen in Figure 9 Taking a median value for the slip during a 250 msec fixation of approximately 0.8 deg, the added retinal velocity would be 3.2 deg/sec at the fovea. We computed how retinal slip of this magnitude would affect speed distributions across the retina. The flow fields in the two cases (perfect stabilization and 0.8 deg of slip) are shown on the left of the figure, and the speed distributions are shown on the right. The bottom plot shows a heat map of the difference. The added slip also has the effect of slightly shifting structure in the motion pattern upwards, by however far from the fovea the eccentric location with the same speed as the slip is. When considering the average signal this shifts the zero point upwards to 4 deg of eccentricity for 4 deg/s of retinal slip. The change in speed is quite small, and the other structural features of the signal are conserved (including the radially asymmetric eccentricity versus speed gradient and the variation with gaze angle).

**Figure 9:**
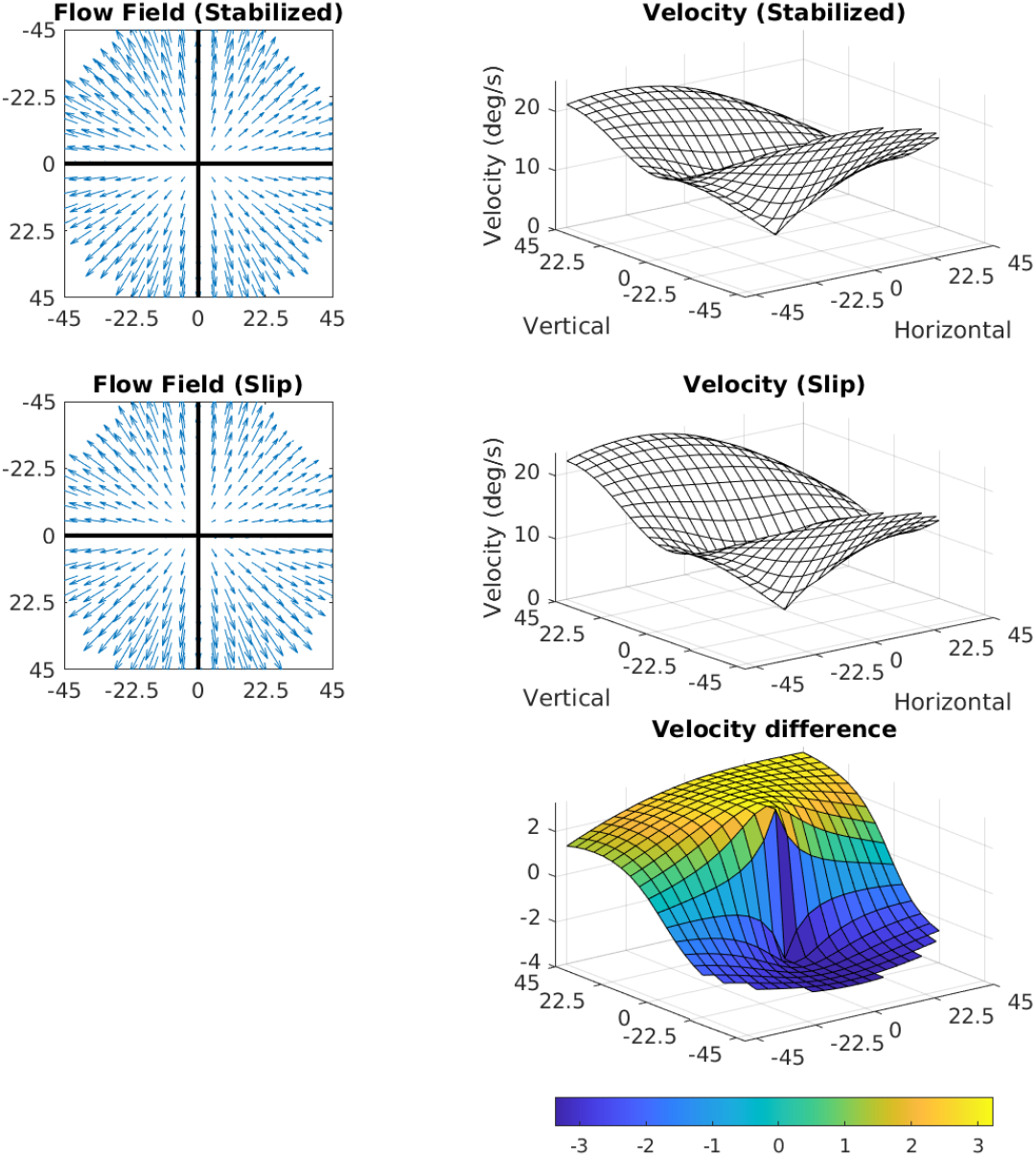
Effects of 3.2 deg/s slip on velocity. The median slip of approximately 0.8 deg during a 250 msec fixation would lead to a retinal velocity of 3.2 deg/sec at the fovea. The figure shows how retinal slip of this magnitude would affect speed distributions across the retina. The flow fields in the two cases (perfect stabilization and 0.8 deg of slip) are shown on the left, and the speed distributions are shown on the right. The bottom plot shows a heat map of the difference.

### Motion generated by saccades

We have thus far considered only the motion patterns generated during the periods of stable gaze, since this is the period when useful visual information is extracted from the image. For completeness, we also examine the retinal motion patterns generated by saccades, since this motion is incident on the retina, and it is not entirely clear how these signals are dealt with in the cortical hierarchy.

### Direction histogram

First we show the distribution of saccade velocities and directions in Figure 10. Saccade velocities are generally less than 150 deg/s as might be expected from the high frequency of small movements to upcoming foothold locations. The higher velocities are likely generated as the walkers saccade from near to far, and vice versa. The direction distribution shows the over-representation of upward and downward saccades. The upward saccades most likely reflect gaze changes towards new foothold further along the path. Some of the downward saccades are from nearby locations to closer ones when more information is needed for stepping.

**Figure 10:**
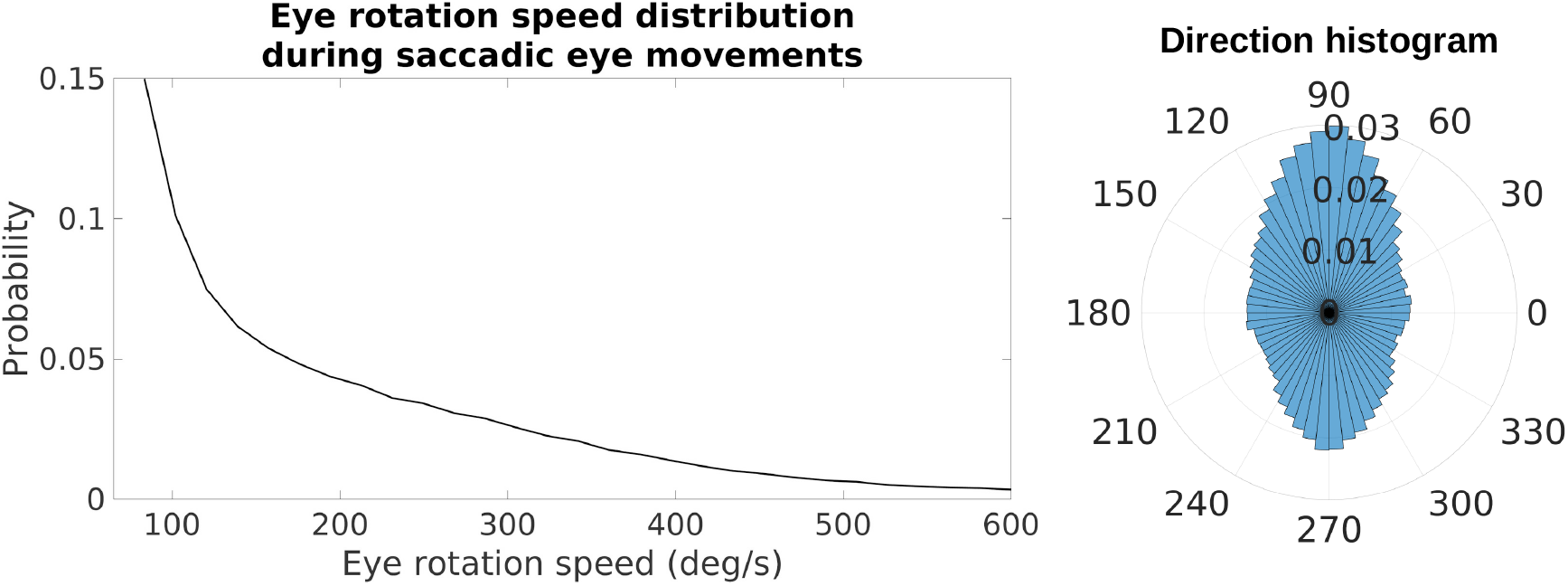
Speed and direction distributions for saccades. The distribution of angular velocities of the eye during saccadic eye movements is shown on the left. On the right is a 2D histogram of vertical and horizontal components of the saccades.

The effect of saccades on retinal image velocities is shown in Figure 11. The speed of the saccade adds to an instantaneous motion input, and shifts motion directions in the opposite direction of the saccade. The Figure shows retinal speed distributions at the 5 locations across the visual field shown in the previous Figures. The top panel shows the distributions previously described, for the periods of stabilization. The middle panel shows retinal speed distributions for the combined data. This shows that the speeds added by the saccades add a high speed lobe to the data without affecting the pattern during periods of stabilization to any great extent. The lower peak resulting from the saccades results from the fact that saccades are mostly of short duration and so account for a lesser fraction of the total data. As expected, the saccades contribute high speeds to the retinal image motion that are largely outside the domain of the retinal speeds resulting from self-motion in the presence of stabilization.

**Figure 11:**
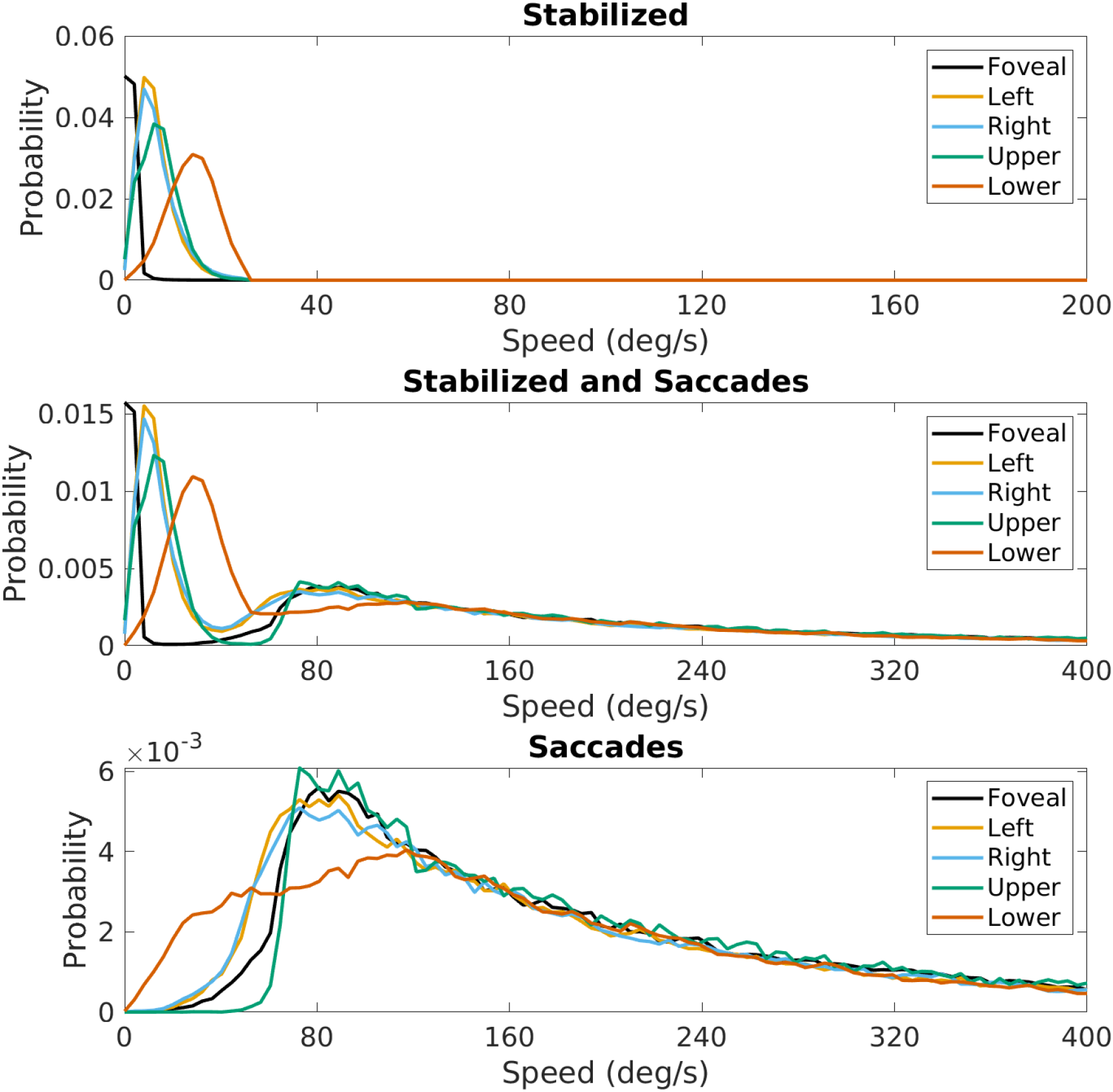
Effect of saccades on speed distributions. Speed distributions at the 5 locations in the visual field shown in Figure 4. The top panel is re-plotted from Figure 4, the middle panel shows these distributions with the saccades added, and the bottom panel shows the contribution from the saccades alone.

